# Genomic Specificity of Anti-TCR mAbs determined by single-cell RNAseq

**DOI:** 10.1101/2024.12.20.629462

**Authors:** Ian Magill, Val Piekarsa, Sofia Kossida, Josh Croteau, Amy Lee, Robert Balderas, David Zemmour, Christophe Benoist, the immgenT Project

## Abstract

T cells play a pivotal role in the immune system, relying on their somatically rearranged T cell receptor (TCR) to recognize peptide-MHC complexes. A comprehensive and extensively used set of monoclonal antibodies (mAbs) against TCR Variable regions was generated in the previous century. The separate identification of mAb-specific TCR-V proteins and *TRV* genes has resulted in multiple nomenclatures, making their relationships unclear. To formally re-establish this link and determine patterns of reactivity within *TRV* subfamilies, we sorted T cells positive for any one of a panel of 22 anti-V mAbs and determined their *TRV* genes by single-cell TCRseq. RNAseq data revealed consistently higher expression of repeated elements from the ERV1-family LTR RLTR6Mm (mapping to *Gm20400*) in cells utilizing *TRBV* segments encoded within a 66kb genomic region between *TRBV23* and *TRBV30. O*ur findings provide a comprehensive resource for anti-TCR mAb specificity and insight into V-gene usage biases and T cell function.

## INTRODUCTION

T cells are central players of the adaptive immune system, orchestrating cell-mediated and antibody-mediated immune responses to protect jawed vertebrate organisms from disease, while enforcing tolerance to self. Through their unique membrane-bound T Cell Receptor (TCR), T cells continuously surveil the presence of self or foreign peptides presented by major histocompatibility complex (MHC) molecules on the surface of antigen-presenting cells. The TCR is a heterodimeric, disulfide-linked glycoprotein, composed of an alpha and a beta chain (alternatively, gamma and delta). As for immunoglobulins, TCR-encoding gene segments are generated through somatic recombination of genomic DNA segments during T cell differentiation: one of an array of variable regions (Vα, Vβ, encoded by *TRAV* and *TRBV* genomic segments) is randomly joined by RAG-mediated recombination to a joining region (Jα, Jβ, encoded by *TRAJ* or *TRBJ* genes), with an additional D element (Dβ) for beta chains. Transcripts from these rearranged elements are joined by splicing onto constant (Cα, Cβ) elements to form the complete TCR mRNAs. This combinatorial assembly, along with imprecise joining processes (base deletions, addition of random N nucleotides), yields a theoretical αβT repertoire of up to 10^15^ unique TCRs in mice and humans. However, the actual repertoire is considerably smaller (1, 2), constrained by the need to conform to MHC molecules.

In the 1980s and 1990s, around the cloning and elucidation of TCR genomics (3), monoclonal antibodies (mAbs) specific for one or a few Vα or Vβ regions were generated (4–25). Such mAbs were generated by several laboratories, through broad immunization with whole T cells across xenogeneic barriers (e.g., rat anti-mouse, hamster anti-mouse) or immunization against specific T cell clones or recombinant proteins. The majority of these mAbs targeted Vβ chains, fewer were specific for Vα chains. These mAbs, initially developed in academic labs, were later compiled into commercially-available panels. These reagents became valuable and versatile tools to profile TCR diversity, track the positive and negative selection events that occur during repertoire selection, monitor transgenically-encoded TCRs, and chart clonal expansion in disease models (8, 10, 26–29).

One of the longstanding issues has been to connect the specificity of these reagents directed against TCR variable regions with the genomic elements encoding those regions. Initially, this connection was established by PCR amplification and cDNA sequencing. However, as the number of identified V regions grew, and as successive generations of genetic nomenclature for V-encoding genomic segments diverged from the original Vα and Vβ denominations (30–32), the linkage between reagents and genomic elements became increasingly less certain. In the late 1990s, the International ImmunoGeneTics information system (IMGT) group formalized the genetic nomenclature of mouse and human *TRV* genes, organizing them according to chromosomal position into a nomenclature system that is currently the accepted standard (33). The IMGT group also performed a retrospective sequence analysis, matching reference sequences to initial publications to infer the correspondence between *TRAV* and *TRBV* genes and anti-V region mAbs (33, 34) This inference was not verified by experimental validation, however, which was particularly an issue for families of closely-related V regions.

In certain cases, the phenotype of a T cell is influenced by the TCR it expresses. For example, superantigen effects during thymic selection can bias cells utilizing specific TCR V genes towards either CD4 or CD8 lineages (35). T cells which recognize nonclassical MHC proteins, such as CD1d and MR1 during thymic development are driven into distinct cell fates, including NKT (36) and MAIT (37) cells, respectively. This differentiation can occur with either germline-encoded invariant TCRs, as seen in type I NKT cells, or with more diverse TCRs that recognize the nonclassical MHC protein, as observed in type II NKT cells (38–40). In γδ T cells, certain combinations of γ and δ variable chains are known to be strongly associated with distinct phenotypes and tissue localizations, such as TRGV5+TRDV1+ dendritic epidermal T cells in mice (41).

The advent of single-cell RNA sequencing has enabled the high-resolution profiling of individual T cells, simultaneously capturing mRNA expression and the sequence of paired α and βTCR components in individual cells (42, 43). This technology has been applied in several large-scale “Atlas” studies, notably in the “immgenT” project, a large multi-center consortium which aims to establish the whole range of states that T cells can adopt in the mouse, at both the RNA and protein levels, alongside the TCRs that are found in such cell states (44). In the immgenT context, we revisited and sought to definitively establish the relationship between the protein V-regions recognized by anti-TCR monoclonal antibodies and the *TRAV* and *TRBV* genetic elements in IMGT-standard nomenclature.

## RESULTS AND DISCUSSION

### Ascribing anti-V mAb reactivity to TRAV and TRBV genes

To directly relate anti-V reactivity and *TRAV/TRBV* regions, we sorted by flow cytometry CD3+ T cells that bound any one of a large panel of anti-mouse V region reagents and analyzed the sorted cell pools by combined single-cell RNA sequencing (scRNA-seq) and TCR sequencing (TCR-seq, both α and β chains). Such data should show the actual combinations of Vα and Vβ regions employed by mAb-bound cells, reveal possible cross-reactivities, and identify transcriptional bias at the mRNA level. Cells labeled with the different mAbs (Fig. 1A) were tagged with DNA barcoded “hashtags” and then pooled for profiling in the same scRNA-seq run. We used T cells (spleen and lymph node cells combined) from a single C57BL/6J mouse obtained from the Jackson Laboratory for optimal genetic traceability.

**Fig. 1.**
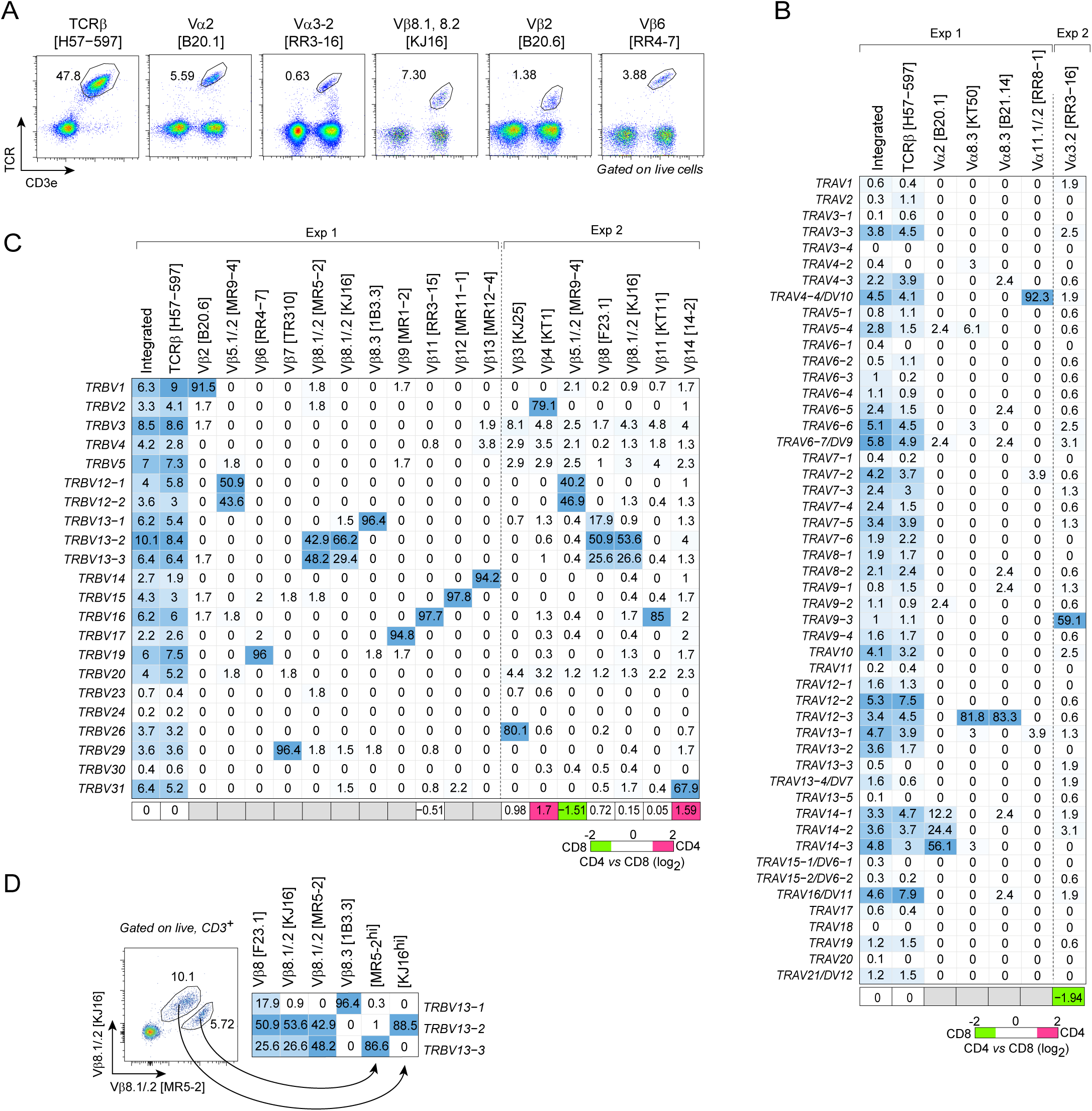
TCR V gene mapping of anti-TCR-V monoclonal antibodies. (A) Flow cytometric profiles of 6 representative mAbs gated on live cells from pooled secondary lymphoid organs of C57Bl/6 mice, as used for sorting for scTCR-seq. (B, C) Frequencies of cells expressing particular *TRAV* (panel B) or *TRBV* (panel C) among cells sorted with one of the mAbs shown at top, and analyzed by scTCR-seq. Frequencies among cells sorted with the pan-TCRβ mAb H57-597 are shown at left, along with frequencies in a broader set of C57Bl/6 T cells from baseline immgenT datasets (“Integrated”). The bottom row shows each mAb’s preferential usage in CD4+ (pink) or CD8+ (green) T cells (as log2 fold change ratio), normalized to pan-TCRβ mAb CD4/CD8 ratio (1.6, **Supplementary Table S3**). Columns with n<100 were blocked out in grey. (D) T cells stained with both anti-Vβ8.1/.2 mAbs KJ16::FITC and MR5-2::PE were sorted into KJ16hi and MR5-2hi, and their *TRBV* usage determined by scTCR-seq as above (data for each anti-Vβ8 individually are reproduced for reference).

These experiments were conducted in collaboration with the two main suppliers of such reagents, BioLegend and Becton Dickinson (Santa Cruz was also contacted, but chose not to participate). General-catalog mAbs, conjugated to either PE or FITC fluorophores, were used for labeling and sorting (listed in **Supplementary Table S1**, examples in **Fig. 1A**). Two independent experiments were performed. A few mAb clones were analyzed twice, with equivalent results. Altogether, the data encompass 17 anti-Vβ and 5 anti-Vα mAbs (the H57-597 mAb was included as a pan-TCRβ control). We used the 10x Genomics 5’v2 with Feature Barcoding protocol for encapsulation and construction of the TCR and mRNA single-cell libraries, with primary data processing using CellRanger (10X Genomics). TCRα and TCRβ contigs generated by CellRanger were processed to determine V, J and N components using IMGT/V-QUEST software (45, 46), and the most updated reference library for C57BL/6J *TRAV* and *TRBV* regions (release 202430-2).

The data obtained ranged from 32 to 462 QC-passing cells for each of the 22 mAbs tested. The *TRAV* and *TRBV* assignments for individual cells in each pool are represented as heatmaps in **Figs. 1B** and **1C** (as percentage of the cells sorted with one particular mAb, with raw numbers listed in **Supplementary Table. S2**). Overall, >90% of cells bound by each mAb corresponded to a specific TCR V gene segment, or a family thereof. For instance, >95% of cells sorted with anti-Vβ6, 7, 9, 11, 12 or 13 all used *TRBV19, 29, 17, 16, 15,* and *14*, respectively. We observed a slight background of low-level cross-contamination, stronger in experiment 2, which we ascribe to imperfect cell sorting: given the number of sorts that needed to be performed in a short timeframe, we only used single-sorting, which yielded purity rates of 90-96% in resorting experiments – not unexpected, given the starting frequencies of positive cells in the 3-15% range. Some mAbs bound to proteins of several members of a family: anti-Vβ5.1/.2 (clone MR9-4 pulled down equal proportions of *TRBV12-1* and *TRBV12-2* positive cells, and the anti-Vα2 (clone B20.1) mAb selected cells expressing any one of the whole *TRAV14* family (*TRAV14-1, 14-2* and *14-3*). In contrast, some mAbs were specific for only one member of a family (e.g. anti-Vα3.2 (clone RR3-16) only identifying *TRAV9-3*-expressing cells, but not those using *TRAV9-1*, *9-2* or *9-4*). The four mAbs specific for the Vβ8/*TRBV13* family (MR5-2, KJ16, F23.1 and 1B3.3) showed partially overlapping patterns of reactivity, as expected from prior reports (***4, 5, 7, 25***). To clarify these relationships, we stained CD3+ cells with both KJ16 and MR5-2, which revealed distinct populations of cells preferentially bound by MR5-2 or KJ16 (**Fig. 1D**). Each of these two populations express either *TRBV13-2* or *TRBV13-3*, confirming the quantitatively preferential cross-reactivity of KJ16 and MR5-2 for *TRBV13-2* or *TRBV13-3*, respectively. In contrast, 1B3.3 proved uniquely specific for *TRBV13-1*, while all three members were bound by F23.1

This data formally establishes the *TRBV* and *TRAV* products recognized by these mAbs, and should serve as useful reference, compiled in **Table 1**. In fairness, the inferences that had been made earlier (33, 47) correctly predicted many of the TRBV relationships established here.

**Table 1.** Anti TCR-V mAb and TRV Gene correspondence.

| Clone | Reference | Protein Name<br>(Barth, 1985) | IMGT Gene Symbol | mAb Does Not Bind |
| --- | --- | --- | --- | --- |
| B20.6 | (Necker et al., 1991) | Vb2 | *TRBV1* |  |
| KJ25 | (Pullen et al., 1988) | Vb3 | *TRBV26* |  |
| KT4 | (Tomonari et al., 1990) | Vb4 | *TRBV2* |  |
| MR9-4 | (Kanagawa et al., 1991) | Vb5.1, 5.2 | *TRBV12-1, TRBV12-2* |  |
| RR4-7 | (Kanagawa et al., 1989) | Vb6 | *TRBV19* |  |
| TR310 | (Okada et al., 1990) | Vb7 | *TRBV29* |  |
| KJ16-1332 | (Haskins et al., 1984) | Vb8.1, 8.2 | *TRBV13-2 > TRBV13-3* | *TRBV13-1* |
| MR5-2 | (Kanagawa, 1988) | Vb8.1, 8.2 | *TRBV13-3 > TRBV13-2* | *TRBV13-1* |
| 1B3.3 | (Kersh et al., 1998) | Vb8.3 | *TRBV13-1* | *TRBV13-2, TRBV13-3* |
| F23.1 | (Staerz et al., 1985) | Vb8 | *TRBV13-1, TRBV13-2, TRBV13-3* |  |
| MR10-2 | (Utsunomiya et al., 1991) | Vb9 | *TRBV17* |  |
| KT11 | (Tomonari and Lovering, 1988) | Vb11 | *TRBV16* |  |
| RR3-15 | (Bill et al., 1989) | Vb11 | *TRBV16* |  |
| MR11-1 | (Abe et al., 1991) | Vb12 | *TRBV15* |  |
| MR12-4 | (Zaller et al., 1990) | Vb13 | *TRBV14* |  |
| 14-2 | (Liao et al., 1989) | Vb14 | *TRBV31* |  |
| B20.1 | (Pircher et al., 1992) | Va2 | *TRAV14* |  |
| RR3-16 | (Utsunomiya et al., 1989) | Va3.2 | *TRAV9N-3* | *TRAV9-1, TRAV9-2, TRAV9-4* |
| KT50 | (Tomonari et al., 1989) | Va8.3 | *TRAV12-3* | *TRAV12-1, TRAV12-2* |
| B21.14 | (Necker et al., 1991) | Va8.3 | *TRAV12-3* | *TRAV12-1, TRAV12-2* |
| RR8-1 | (Jameson et al., 1991) | Va11.1, 11.2 | *TRAV4-4/DV10* | *TRAV4-1, TRAV4-2, TRAV4-3* |

Several earlier studies showed that individual Vα or Vβ chains are preferentially utilized by CD4+ or CD8+ T cells, denoting a preferential interaction with MHC-II or MHC-I molecules (11, 48–50). Because total CD3-positive cells were used in the present analyses, it was possible to determine V gene usage biases between CD4+ and CD8+ T cells (the ratio observed between CD4+ and CD8+ frequencies were normalized to that in cells sorted with the pan-TCRb (H57-597) mAb. (**Fig. 1B,C, Supplementary Table S3**). The results are mostly consistent with previous reports, such as *TRAV4-4/DV10* showing skewed expression towards CD8+ subset (14), *TRBV2* and *TRBV31* being preferred in CD4+ subsets (11, 16), while several others similarly having biased CD4/CD8 ratios.

These experiments also allowed us to ask whether the binding of a mAb to its Vβ target could be prevented by particular Jβ or Cβ elements, or by pairing to specific Vα chains, leading to cells displaying the target Vβ but not actually bound by the cognate mAb (51). To test this, we combined the full anti-Vβ mAb panel (PE- or FITC-conjugated), and sorted the PE- and FITC-negative population of CD3+, TCRγ/δ-T cells (**Fig. 2A**). Among these 2,856 negative cells, only the “untargeted” *TRBV* were found (with exception of a few *TRBV1*+ cells, likely because the anti-Vβ2 lot used yielded low fluorescence – see **Fig. 1A**). Thus, other than this exception, the anti-Vβ mAbs effectively recognized all cells expressing their corresponding targets, with no loss of staining due to interference by particular Cβ, Jβ or Vα (we cannot rule out quantitative effects, however). To confirm this conclusion, we compared the frequencies of TCR V gene usage obtained through mAb staining and those derived from scTCR-seq, in larger numbers of cells in the immgenT data, which proved highly concordant (**Fig. 2B**).

**Fig. 2.**
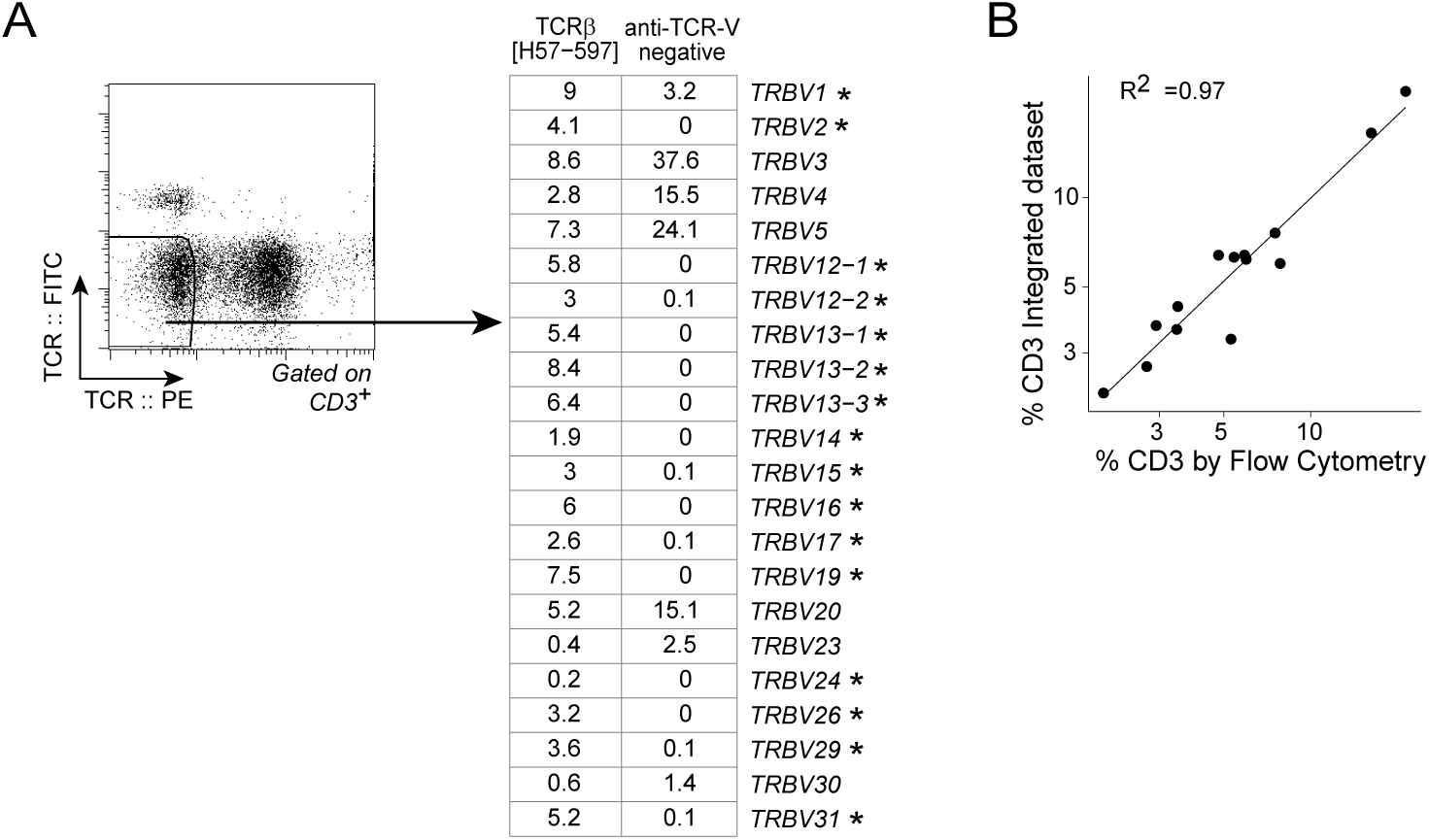
mAbs capture all Vβ-expressing cells. (A) scTCR-seq determination of *TRBV* gene usage among cells sorted as negative for the entire cocktail of anti-Vβ mAbs (left). Right: *TRBV* frequencies among 2856 non-staining cells in the scTCR-seq dataset. Frequencies among H57-597 positive cells are shown as reference. (B) Comparison of *TRBV* usage as determined by flow-cytometry after staining with the mAbs of Table S1 against by scTCR-seq.

### A link between V region usage and cell phenotype ?

The design of this experiment also lent itself to investigating relationships between the V region expressed by a cell and its phenotype as revealed by transcriptome profiles. The sorting of T cells with the anti-TCR mAbs corrected for the uneven utilization of TCR V genes, and captured more cells utilizing a given V segment than if relying on the TCR-seq output of total T cells. For robustness, this analysis was restricted to samples with higher cell numbers (> 200) and was focused on CD4+ T cells to circumvent differential V gene usage in CD4+ and CD8+ T cells from obscuring more subtle differences. Displaying the whole CD4+ T cell population by Uniform Manifold Approximation and Projection (UMAP) showed the usual partition between naïve and effector/memory poles (**Fig. 3A**). Within this framework, cells sorted by usage of particular V regions distributed in all quadrants of the UMAP (**Fig. S1A**). We then performed differential cell density analysis, which showed that cells expressing one particular V region only showed scattered local maxima, similar to fluctuations observed with random sampling of similarly-sized sets of cells, and not particularly concentrated in the naïve or effector/memory regions of the UMAP. space (**Fig. 3B**).

**Fig. 3.**
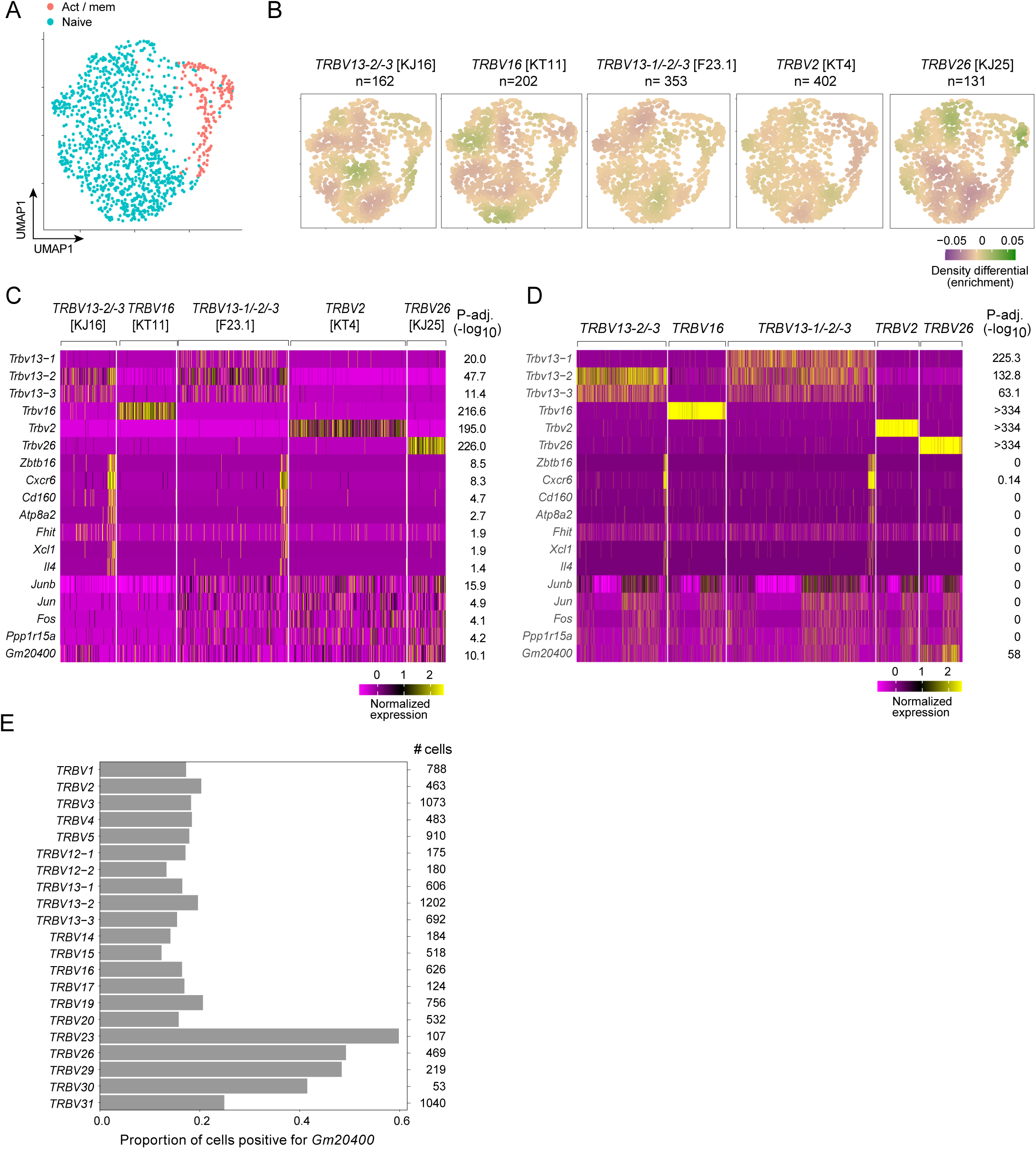
Variation in transcriptional phenotype according to V region usage. (A) UMAP representation of CD4+ T cells in the dataset, color-coded as naïve of activated/memory according to signature gene expression. (B) Cell density plots of CD4+ T cells sorted as positive for the mAb shown, relative to all CD4+ T cells in the dataset. (C) Heatmap of differentially expressed genes between cells sorted with different anti-Vβ mAbs, (D) Heatmap showing the expression of the same genes among all CD4+ T cells found to utilize the same *TRBV* regions in the broader immgenT dataset (4,187 cells altogether in this panel). (E) Proportion of cells scoring as positive for the Gm20400 lnc RNA among cells of the larger integrated dataset, according to their *TRBV* usage (total number of cells for each shown at right).

Differential gene expression analysis between the same sets of cells brought up few differentially expressed genes (**Fig. 3C**), which fell into several categories. First, and comfortingly, the corresponding *TRBV* genes (top 5 rows). Second, a set of seven transcripts known to be over-expressed in iNKT cells, over-represented in Vβ8+ cells sorted with KJ16 and F23.1, including: *Zbtb16* (the master regulator of NKT cells), *Cxcr6, Cd160, Atp8a2, Fhit, Xcl1,* and *Il4.* This enrichment is understandable because in iNKT cells the invariant Vα chain is preferentially paired with a Vβ8/*TRBV13* products. Third, AP-1 related genes (e.g. its components *Junb, Jun,* and *Fos* and a downstream AP-1 target *Ppp1r15a)* were over-represented in *TRBV2* [KT4], *TRBV13-1/-2/-3* [F23.1] and *TRBV26* [KJ25] cells. Finally, and perhaps most intriguingly, cells positive for *TRBV26* [KJ25] were enriched in transcripts from an endogenous retroviral element, RLTR6Mm (mapping in many genomic locations but most homologously to an exon of the long noncoding RNA (lnc) *Gm20400,* located on the same chromosome 6 as the *TRBV* cluster, but 100 Mb downstream (142 Mb vs 41 Mb). This repeated transcript itself includes a duplication of the 35 bp 5’TCTTTCATTTGGTGCATTGGCCGGGAATTCGAGAA3’ motif.

The same patterns of differential gene expression were found in CD8+ cells from the same dataset (Fig. S2B). We then asked whether they could be replicated in cells expressing the same TCR V regions, but identified without use of antibodies within larger sets of CD4+ spleen and lymph node cells from unmanipulated B6 mice analyzed in the immgenT program. From 11975 total CD4+ T cells, we filtered those expressing the same Vβ regions as in the analyses of Fig. 3A-C, and assessed the differential expression of the same genes (**Fig. 3D**). Differential representation of both the iNKT cluster and the AP-1 transcripts largely vanished. The loss of significance for iNKT transcripts was due to a lower representation of iNKT cells relative to the initial experiment, and we interpret the discordance in AP-1 representation as likely reflecting an experimental artefact linked to mAb binding during our experiment, especially since AP-1 activation is part of the typical “stress signature” in scRNAseq datasets (although it is not clear why some mAbs would behave differently from others). Interestingly, expression of the lnc *Gm20400* was again associated with the usage of *TRBV26*. (Fig. 3D, bottom row). Broader examination of all cells in this immgenT dataset revealed that high expression of RLTR6-Mm ERV transcripts assigned to *Gm20400* were found in cells employing one of a restricted set of *TRBV*, including *TRBV23*, *TRBV26*, *TRBV29* and *TRBV30*. (**Fig. 3E**). Interestingly, these *TRBV*s are all encoded within a 66kb stretch of the large *TRBV* complex. It is unclear at this juncture why and how the transcriptional activity of an endogenous retroviral element correlates with the frequency of rearrangement of a subset of *TRBV* regions (no other *TRVB* was associated with overexpression of other repeated elements). There are 251 copies of RLTR6-Mm in the mouse genome (52) and one can speculate that 3D folding of the genome that allows rearrangements of the *TRBV23*-*TRBV30* interval may favor transcription at one of these repeats.

In conclusion, this study establishes a direct link between anti-V region mAb reactivity and *TRAV/TRBV* gene usage, providing a comprehensive and experimentally validated reference for these reagents, confirming their high specificity and the absence of interference by other TCR elements.

## ACKNOWLEDGEMENTS

We thank the Flow Cytometry core at Harvard Medical School for their assistance with FACS. This study was funded by a resource grant from NIH/NIAID to the ImmGen consortium (AI072073).

## DISCLOSURES

RB is an employee of Becton Dickinson, Inc. JC and AL are employees of BioLegend, a Revvity company.

## SUPPLEMENTAL FIGURE LEGENDS

**Fig S1.**
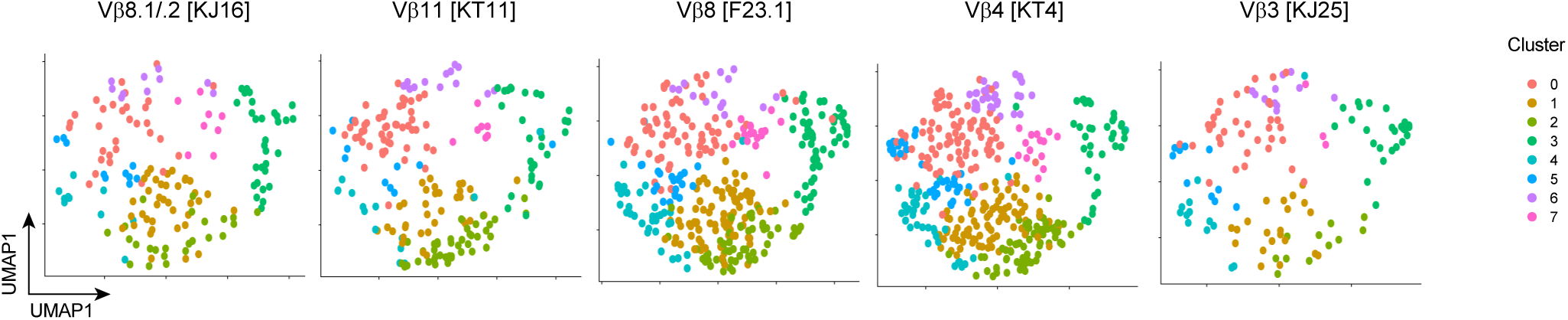
Distribution of *TRBV* positive cells in UMAP space. UMAP representation of CD4+ T cells in the dataset, split by *TRBV* sample.

**Fig S2.**
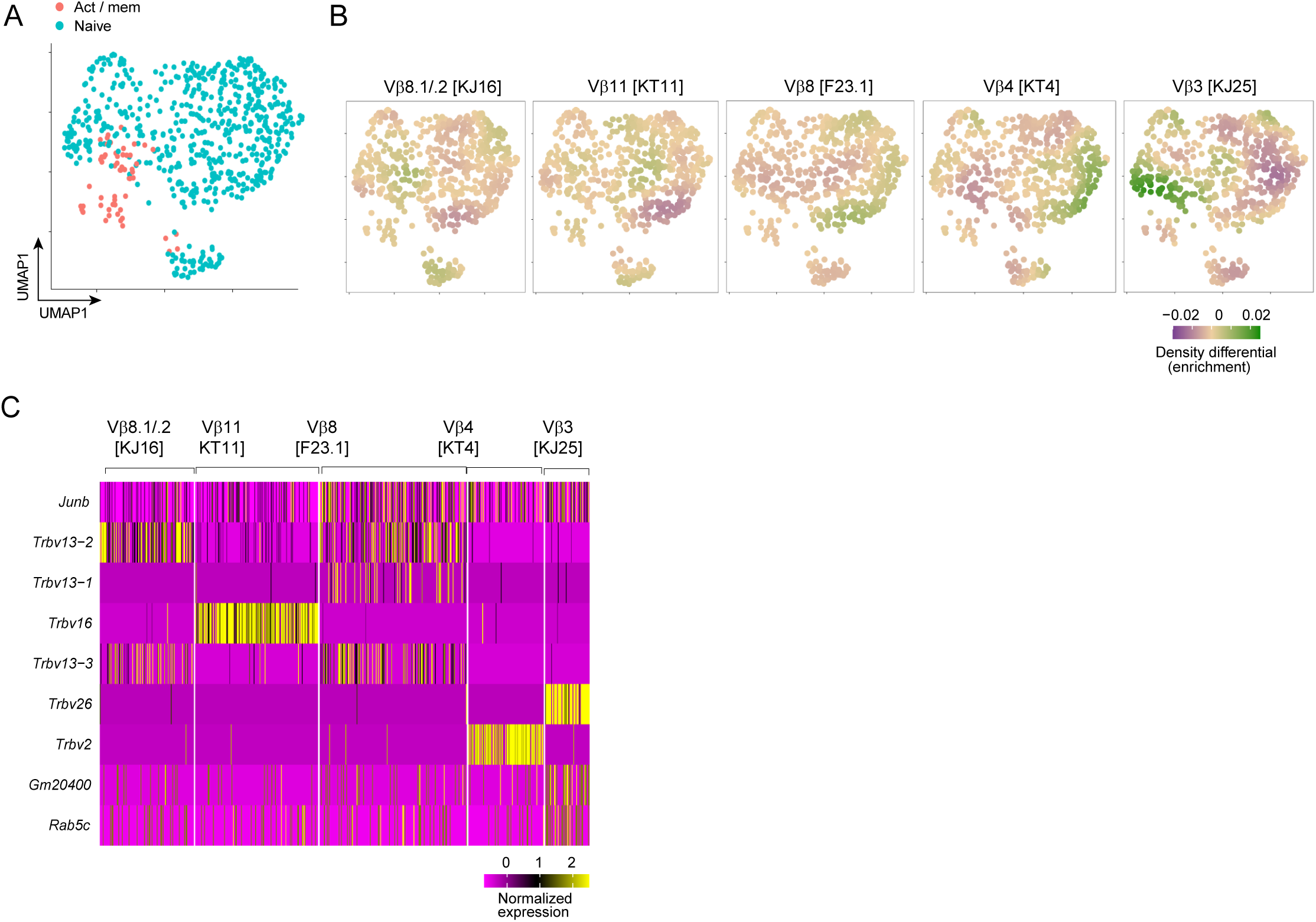
Variation in transcriptional phenotype according to V region usage in CD8+ T cells. (A) UMAP representation of CD8+ T cells in the dataset, color-coded as naïve or activated/memory according to signature gene expression. (B) Cell density plots of CD8+ T cells sorted as positive for the mAb shown, relative to all CD8+ T cells in the dataset. (C) Heatmap of differentially expressed genes between cells sorted with different anti-Vβ mAbs

## METHODS

### Mice

All mice were maintained in accordance with Harvard Medical School’s Animal Care and Use Committee guidelines (IACUC protocol #IS00001257). C57Bl/6J (B6) mice were purchased from the Jackson Laboratory. 7-week male mice were used for all flow cytometry and transcriptomic experiments.

### Cell Isolation and Flow Cytometry

Pooled inguinal, axillary, brachial, mesenteric lymph nodes and spleen were collected into 1.7mL Eppendorf tubes containing 1mL sterile DMEM with 2% newborn calf serum (NBCS), 0.1% Sodium Azide, and 10mM HEPES.

Pooled lymph nodes were mechanically dissociated and filtered through a 40um nylon mesh, then resuspended in buffer. Spleens were processed in the same way, and additionally underwent red blood cell lysis in ACK lysis buffer. The single cell suspension was divided into aliquots for each *TRV* sample and simultaneously stained with both fluorochrome-conjugated monoclonal antibodies to saturation and oligo-conjugated Totalseq-C anti-mouse hashtags (0.5 ug per sample) for 20min in the dark on ice and washed twice (1mL of media, 5 minute spin at 500g). Each *TRV* sample in a given experiment was stained with its associated *TRV* antibody and a unique combination of hashtag antibodies. Each hashtag antibody possesses a unique oligo sequence, ubiquitously binds immune cells, and allows for the assignment of sequenced single cells to their original sample based on the specific oligo-derived reads associated with each cell. In experiment one, a dual-hashing scheme was used where cells were assigned to their original sample based on the presence of specific combinations of hashtag oligos. Dead cells were excluded using 4’, 6-diamidino-2-phenylindole hydrate (DAPI). Cells were sorted once using FACSAria III (BD Biosciences) or MoFlo Astrios (Beckman Coulter Life Sciences) cell sorters. Cells were sorted in relative quantities according to Supplementary Table S1 into a shared collection tube. Flow cytometry data was analyzed in FlowJo v10.10.

### Single Cell RNA-, TCR- and CITE-seq

Single cell RNA-seq was performed in accordance with the immgenT protocol described in (53). Briefly, single cell RNA-seq was performed on the selected cells using the 10x Genomics 5’v2 with Feature Barcoding for Cell Surface Protein and Immune Receptor Mapping protocol, per the manufacturer’s instructions (CG000330). The resulting libraries were sequenced on Illumina Novaseq flow cells.

### scRNAseq processing

Gene and Totalseq-C antibody-derived reads were aligned to mm10(GRCm38) genome and Totalseq-C hashtag DNA barcodes (**Supplementary Table S1**), respectively, using the CellRanger (v8.0.1) software (10x Genomics) (54). Cells and empty droplets were differentiated using the barcodeRanks function in the DropletUtils package. Samples were demultiplexed using the Totalseq-C hashtag reads using the HTODemux function. To demultiplex samples labelled with multiple hashtags, the HTODemux function was repeated twice, with all valid dual combinations of hashtags split across the two repeats. Cells multi-positive for those hashtags were assigned to distinct samples. Cells satisfying any of the following conditions were excluded from the analysis: dead cells with >5% mitochondrial genes; fewer than 471 RNA transcripts; fewer than 500 detected genes; hashtag-negative cells; and hashtag-multipositive cells (except for intentionally dual-hashed samples). Dimensionality reduction, visualization, and cluster analysis was performed using the Seurat v5.1 package.

Differential cell density was calculated using the MASS package. Differential gene expression analysis was done with Seurat’s FindAllMarkers function. For the analysis of the integrated ImmgenT dataset, cells in the *TRBV13* family were grouped to better match the representation in the antibody-sorted samples: cells expressing *TRBV13-2* and *TRBV13-3* were randomly split into two equal populations. Half of the *TRBV13-2* and *TRBV13-3* cells were merged to create the KJ16[Vβ8.1/.2] analogue. The *TRBV13-1* cells and the other half of the *TRBV13-2* and *TRBV13-3* cells were merged to create the F23.1[Vβ8.1/.2/.3] analog.

Reads assigned by CellRanger to *Gm20400* were visualized on the UCSC Genome Browser which showed the assigned reads aligning primarily to a narrow region of exon 2. Homologous sequences were identified using BLAST (Ensembl) or MUSCLE (55). Homologous sequences were screened for interspersed repeats using RepeatMasker (56).

### TCR processing

TCRαβ and TCRβ contigs for each cell were generated by aligning reads to reference genes using CellRanger (v8.0.1) with a custom reference of C57Bl/6 elements. These contigs were then processed using IMGT/V-QUEST (using the same B6 reference library), resulting in tables of parsed V, D, J and N elements for each cell.

